# Retinal organoids derived from rhesus macaque iPSCs undergo accelerated differentiation compared to human stem cells

**DOI:** 10.1101/2021.05.25.445693

**Authors:** Antonio Jacobo Lopez, Sangbae Kim, Xinye Qian, Jeffrey Rogers, J. Timothy Stout, Sara M. Thomasy, Anna La Torre, Rui Chen, Ala Moshiri

## Abstract

**Purpose:** To compare the timing and efficiency of the development of non-human primate (NHP) derived retinal organoids in comparison to those derived from human embryonic stem cells.

**Methods:** Human embryonic stem cells (hESCs) and induced-pluripotent stem cells (rhiPSCs) derived from non-human primates (*Macaca mulatta*) were differentiated into retinal organoids by using an established differentiation protocol. Briefly, embryoid bodies were formed from pluripotent stem cells and induced into a neural lineage with neural induction media with the addition of BMP4. Thereafter, self-formation of optic vesicles was allowed to form in a 2D culture in retinal differentiation media (RDM). Optic vesicles were then manually harvested and cultured in suspension in 3D-RDM media until analysis. Differences in the timing of differentiation and efficiency of retinal organoid development were assessed by light microscopy, electron microscopy, immunocytochemistry, and single-cell transcriptomics.

**Results:** Generation of retinal organoids was achieved from both human and several NHP pluripotent stem cells lines. All rhiPSC lines resulted in retinal differentiation with the formation of optic vesicle-like structures similar to what has been observed in hESC retinal organoids. NHP retinal organoids had laminated structure and were composed of mature retinal cell types including cone and rod photoreceptors. Single cell RNA sequencing was conducted at two time points, which allowed identification of cell types and characterization of developmental trajectory in the developing organoid. Important differences between rhesus and human cells were measured regarding the timing and efficiency of retinal organoid differentiation. While the culture of NHP-derived iPSCs is relatively difficult compared to human stem cells, the generation of retinal organoids is feasible and may be less time consuming due to an intrinsically faster timing of retinal differentiation.

**Conclusions:** Retinal organoids produced from iPSCs derived from Rhesus monkey using established protocols differentiate through the stages of organoid development faster than those derived from human stem cells. The production of NHP retinal organoids may be advantageous to reduce experimental time and cost for basic biology studies in retinogenesis as well as for preclinical trials in NHPs studying retinal allograft transplantation.

## INTRODUCTION

High acuity vision in primates is attributed to the development of the macula lutea (macula), a region with a high density of cone photoreceptors and specialized circuitry. The center of the macula is the fovea centralis (fovea), which is devoid of retinal vasculature and has a characteristic depression, the foveal pit, which includes only tightly packed cone photoreceptors at the exclusion of rods. Many forms of retinal blindness, whether age-related or congenital, ultimately lead to a final common pathway of vision loss due to foveal damage. Inherited retinal disease (IRD), typically associated with single gene mutations, can cause macular dystrophies and cone disorders which harm the fovea. Age-related macular degeneration (AMD) and related conditions can lead to damage to the fovea. Retinal vascular disorders such as diabetic retinopathy and occlusive diseases of the retinal vasculature also lead to foveal damage through a combination of macular ischemia and macular edema. The final common pathway of most forms of retinal blindness is macular damage and loss of foveal cones and their supporting cells. Therefore, there is a need for the advancement of cellular regenerative or replacement technologies for patients who have vision loss secondary to foveal cell death. One promising technology is cellular transplantation of cone photoreceptors to the macula to restore vision. As terminally differentiated cells, photoreceptors do not have the ability to regenerate. The promise of pluripotent stem cells (PSCs) as a source of photoreceptors through *in vitro* differentiation has greatly advanced with studies demonstrating the ability to produce consistent and reproducible 3D retinal organoids^1,2^. These results show promise in the possibility for photoreceptor replacement treatments. Transplantation of photoreceptors derived from human pluripotent stem cells have demonstrated promise in rodents,^3,4,5^ felines,^6^ and non-human primates ^7,8,9^. However, transplantation experiments to the retina of mammals with a macula are relatively few and successful restoration of vision has been limited.^10,11,12^ The translation to clinically relevant treatments relies on the ability to accurately model these therapeutic approaches in a context similar to human patients. While rodents, felids, and canids provide important data on safety and efficacy, they lack true foveal architecture and therefore, they are not a complete model for the complexity of human macular diseases. It is important to demonstrate the safety and efficacy of these treatments in preclinical trials. Perhaps the best models we have for preclinical macular studies are non-human primates. Non-human primates (NHP) are highly genetically, anatomically, physiologically, and behaviorally similar to humans and they represent excellent models for translational research when other animal models do not recapitulate the disease of interest. Rhesus macaques (*Macaca mulatta*) are one of the most commonly used NHPs in biomedical research. We have recently described macaques with inherited retinal disease which may benefit from transplantation of retinal tissue to restore macular function.^13^ Others have also described macaques with macular abnormalities.^14,15,16^ As the number of NHP models of IRDs expand, these resources can be useful to study cell transplantation in the context of pre-existing retinal disease. Examples of allogeneic cell transplantation are limited.^17^ Exploration of transplantation of allogeneic retinal cells is necessary to overcome the barriers to human clinical studies. Therefore, we have sought to differentiate three-dimensional retinal organoids from induced-Pluripotent Stem Cells (iPSCs) derived from Rhesus monkeys for the purpose of transplantation into animals with IRDs.

Similar to human stem cells, Rhesus macaque iPSCs (rhiPSC) are characteristically indistinguishable from human iPSCs. A previous study in cynomolgus macaque (*Macaca fascicularis)* derived embryonic stem cells demonstrated that the production of retinal cell types from NHP iPSCs is possible using a two-dimensional differentiation protocol.^18^ However, while rhiPSCs have been used for the derivation of blood products^19^ and cardiac cells,^20,21,22^ little is known about the production three-dimensional retinal organoids from rhiPSCs. In this study, we demonstrate the ability of rhiPSCs to produce 3D retinal organoids using a composite established retinal differentiation protocol.^23,24,25^ We characterize the development of rhiPSC derived 3D retinal organoids and their composition with immunocytochemistry and single-cell next-generation sequencing. To determine the reproducibility of the differentiation protocol, we use three rhesus iPSCs: two established rhiPSC89^26^ and rhiPSC90^27^; and a novel line (from the lab of James Thomson), rhiPSC2431, that is in a defined and xeno-free culture system. Our results demonstrate that rhiPSC retinal organoids follow an abbreviated, but similar development to human iPSC-derived retinal organoids which is consistent with the reduced gestational period of rhesus macaques.

## RESULTS

### Rhesus iPSCs differentiate into 3D retinal organoids

In this study, we differentiated three different rhiPSC lines using a stepwise retinal differentiation protocol to generate retinal organoids (Figure 1A). Undifferentiated stem cell colonies were cultured in non-adherent conditions to generate embryoid bodies (EBs), which were cultured in neural induction media (NIM) for 7 days. Next, the EBs were plated onto Matrigel-coated plates and grown in 2D from day 7 to 30 at which time, the regions of the culture displaying retinal morphology were selected, lifted, and grown in non-adherent 3D conditions (Figure 1B). The morphology of the rhiPSC-derived cultures developing retinal tissue was very similar to that observed in H9 hESCs. Following the criteria proposed in Caposwki et al (2019), we binned retinal organoids into three stages: stage 1, retinal organoids displaying phase bright appearance; stage 2, retinal organoids displaying phase dark appearance; and stage 3 retinal organoids displaying outer segment protrusions (Figure 1C). During the 2D-retinal differentiation protocol, we noticed that the self-organizing into optic vesicle like structures (OVs) developed earlier and were well formed as early as day 20 (Figure 1B, Day 20). OVs formation was not mutually exclusive to formation of neural rosettes (Lamba et al., 2009a) or horseshoe-like structures described by others (Zhong et al., 2014). OVs continued to develop until day 30, at which point they were manually selected from the plate and resuspended in 3D-RDM media supplemented with retinoic acid (RA) until day 80. Thereafter, the retinal organoids were cultured in 3D-RDM until analysis.

**Figure 1:**
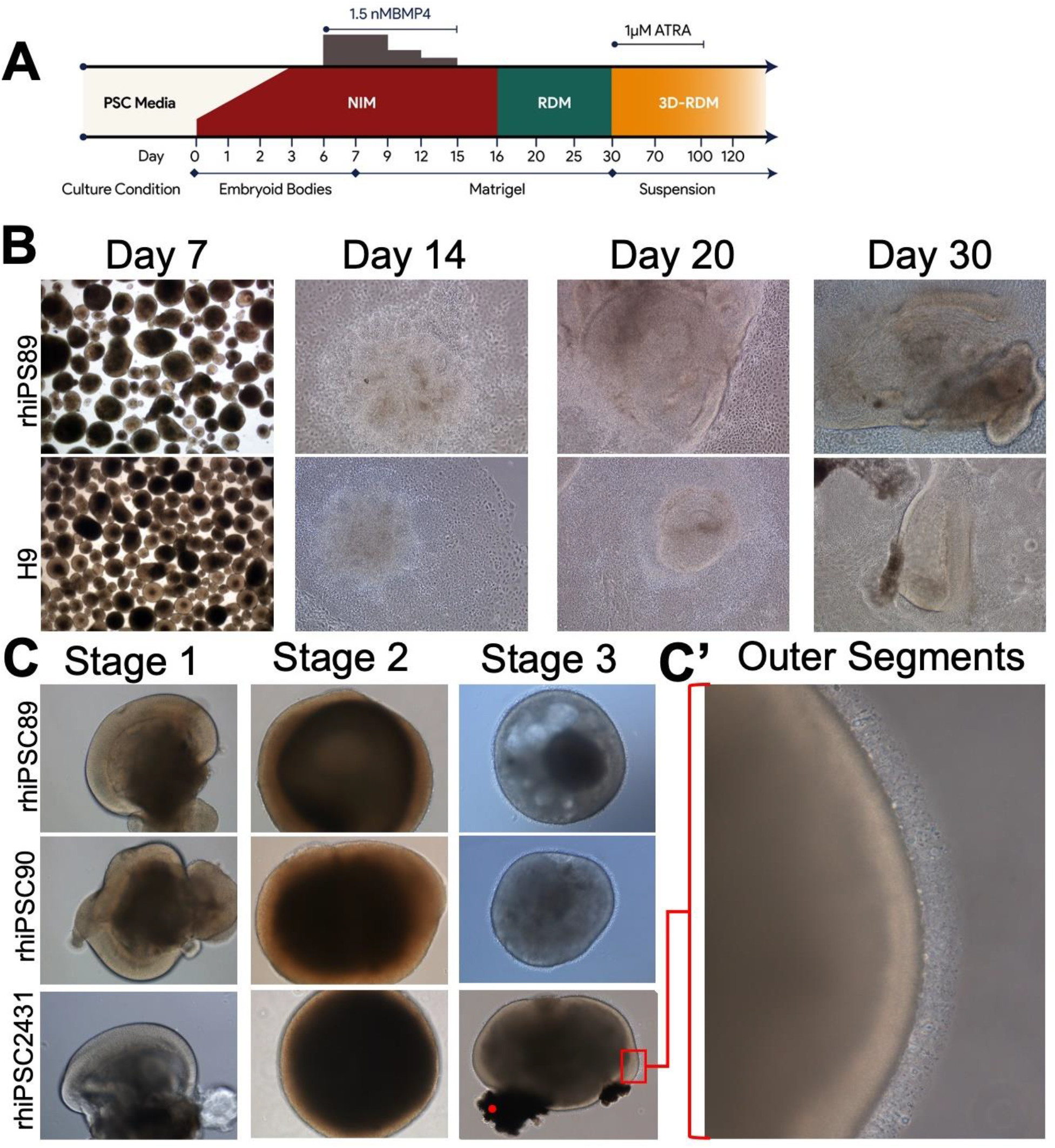
Schematic diagram and phase microscopy of rhesus retinal differentiation protocol. A) Schematic diagram of the differentiation protocol used to produce 3D-rhesus-retinal organoids. B) Representative phase images of the differentiation protocol before plating on Matrigel coated plates (day 7), rosette formation (day 14) and the development of optical vesicle-like structures (day 20 and day 30). C) Representative phase images of rhesus retinal organoids in suspension (stage 1, 44days; stage 2, 80 days; and stage 3, 105days (rhipsc89 and 90) and 125 days (rhiPSC 2431). C’ High magnification image of the marked area demonstrating the hair-like extrusions on the perimeter of the retinal organoids

### Rhesus retinal organoids undergo appropriate morphological and structural differentiation

Stage 1 retinal organoid sections were examined (day 44) with light microscopy (Figure 2A). At this stage, some organoids had a variable number of cells in the inner layers of the organoids. Smaller organoids usually had higher density of cells in their core (Figure 2A’) while larger organoids as they transition to stage 2, developed a hollow core. There was further loss of the inner cells as retinal organoids developed to stage 3. The outer aspect of rhesus retinal organoids had discrete shape and boundaries facing their environment, while the internal boundary was poorly defined (Figure 2A, 2A’). Stage 2 and 3 organoids were similar to one another in that they had a loss of inner cells. However, we also observed a thickening of the outer layer. Similar to hiPSC-derived retinal organoids, stage 3 was defined with the development of outer segment-like morphology (Figure 1C’). To further elucidate whether the structures observed were photoreceptor segments, stage 3 retinal organoids (125 days) were analyzed through transmission electron microscopy (Figure 2B, 2B’, 2C, and 2C’). The photoreceptor protrusions are composed mostly of inner segment-like structures (figure 2B and B’, IS) that are mitochondria rich (2B’, MT). There is also a presence of a loose outer limiting membrane (Figure 2B and B’, OLM). Outer segments (OS) and connecting cilium structures were also observed (C, OS and C’ CC, respectively). These ultrastructural findings demonstrate that stage 3 retinal organoids can develop photoreceptor structures similar to those observed in human retinal organoids.

**Figure 2:**
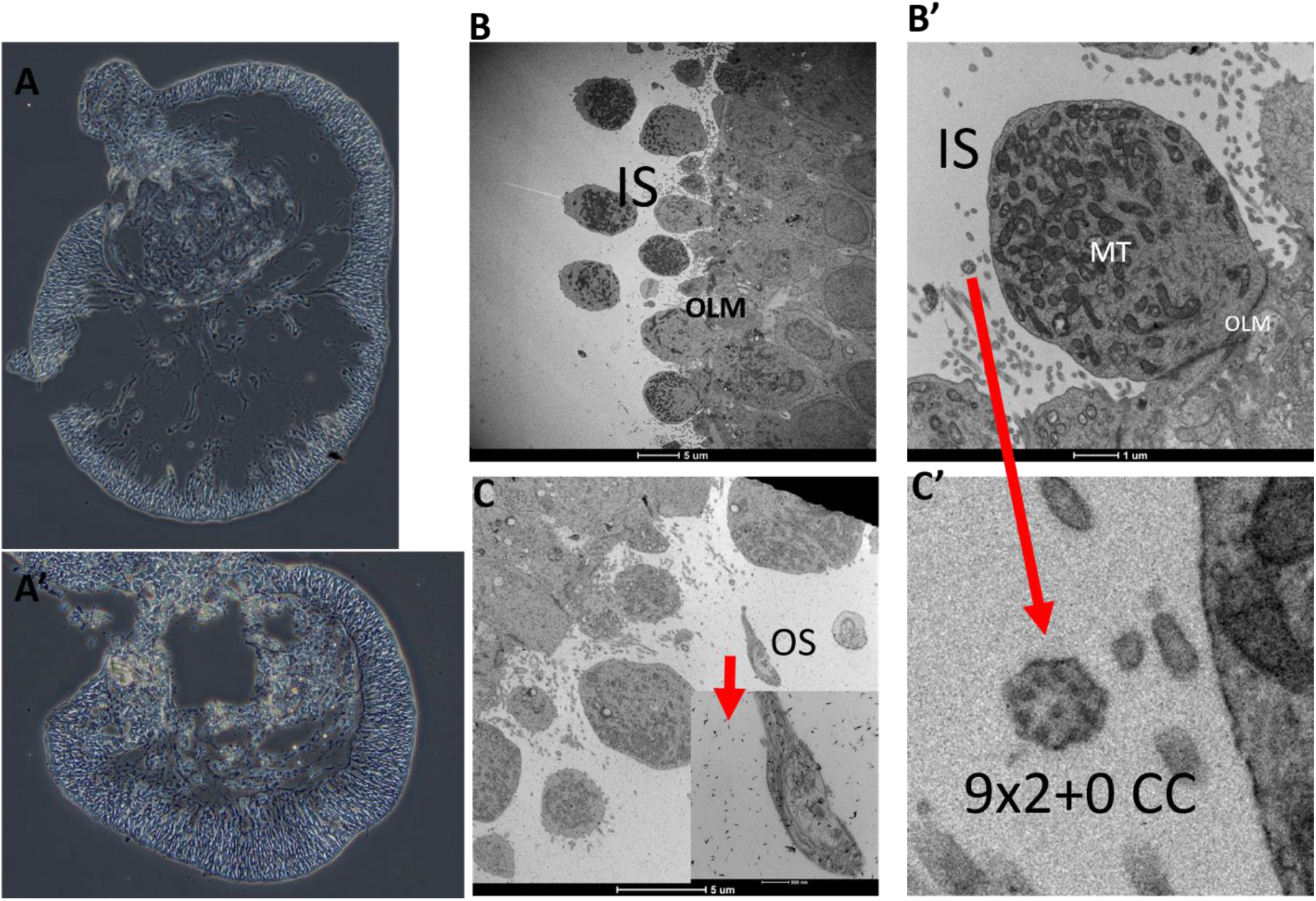
Light microscopy and transmission electron microscopy structural analysis of rhesus retinal organoids. A) Phase image of a 44day old rhesus-retinal organoid. The perimeter of the retinal organoids is cell dense, while internally, cells are sparse. A’) Phase image of a smaller 44 day old rhesus-retinal organoid that contains more internal cells. B) Transmission electron microscopy of the perimeter of stage 3 rhesus retinal organoids (125 days) demonstrating inner segments (IS), an outer limiting membrane (OLM) B’) Inner segments are mitochondria (MT) rich. C) outer segment (OS). C’) Magnification of B’ demonstrating a 9×2+0 structure of connecting cilia (CC).

### Rhesus retinal organoids express retinal cell type-specific markers

Stage 1 retinal organoids are composed of neural retinal progenitors, retinal ganglion cells and early photoreceptors. Immunocytochemistry of stage 1 retinal organoids demonstrate an abundance of eye field transcription factors and retinal progenitor markers such as Pax6, Rx, Lhx2, Otx2, and Chx10 (Figure 3 A, B, F, L and M). In human retinal organoids, early photoreceptors localize to the outer perimeter. Cells expressing both Otx2 and Blimp1 (Figure 3H-K) are committed to photoreceptor fates (Brzezinski, Lamba, & Reh, 2010). By using these fate restriction markers, we were able to determine that as early as stage 1, early photoreceptors (presumably cones) are primarily localized to the outer perimeter of the organoid. Similarly, we also observed double-positive cells expressing both Recoverin and Crx, further indicative of photoreceptors precursors (Figure 3 O-R). Crx/Recoverin positive cells primarily resided in the outer perimeter of the organoids, while other cells lacking these markers resided in the inner layers. However, they were not restricted to the innermost aspect of the organoids. Furthermore, markers of bipolar cell restriction (Otx2+/Chx10+) were also present at this stage (Figure 3L-N).

**Figure 3.**
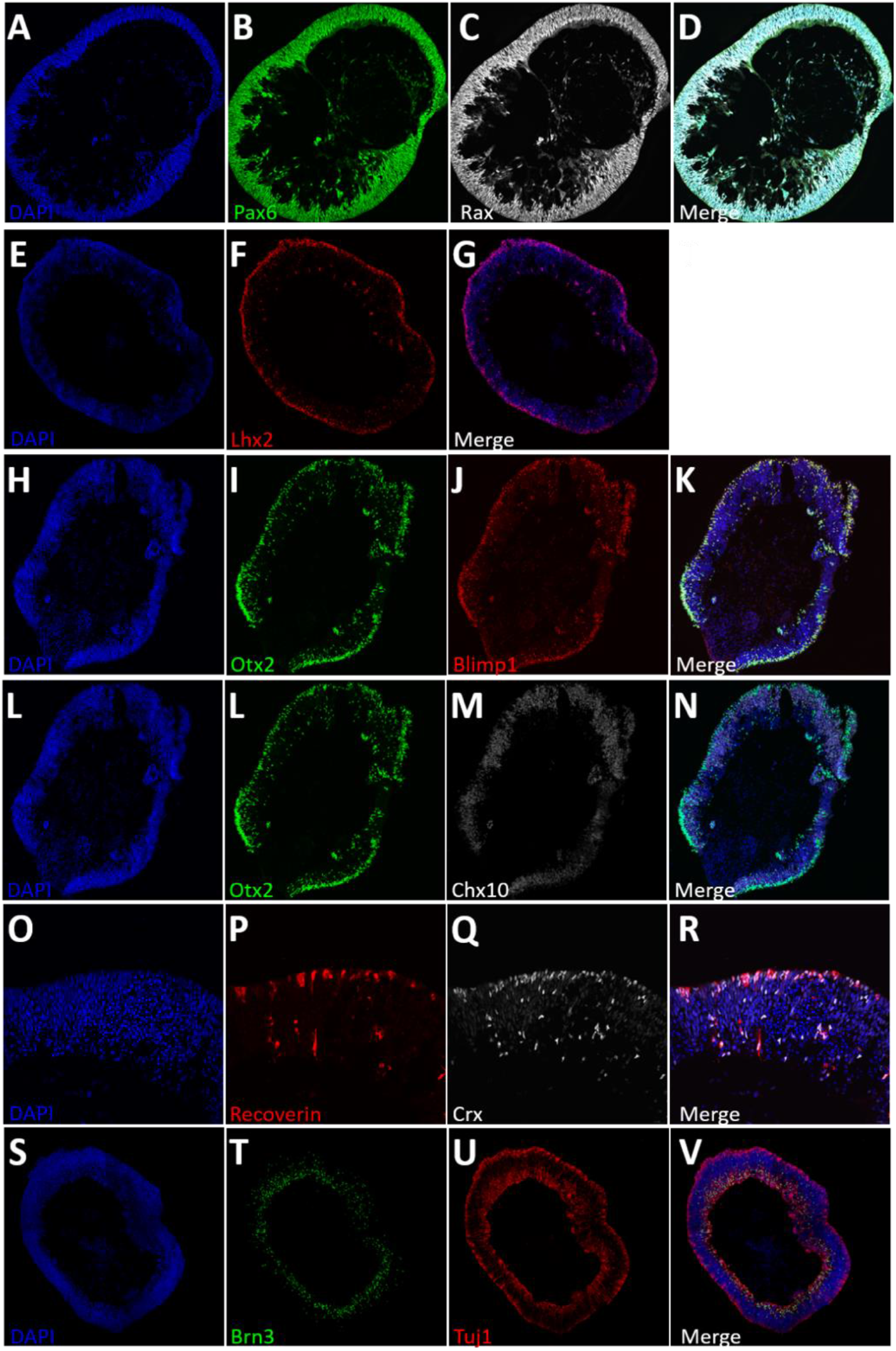
ICC of Stage 1 (44 days): Retinal organoids are composed of neural retinal progenitors, retinal ganglion cells and early photoreceptors. (A, B, F, L and M) Retinal organoids express eye field transcription factors Pax6, Rx, Lhx2, Otx2, and Chx10, respectively. (H-K) Retinal organoids contain Otx2+/Blimp1+ cells, markers of early photoreceptor fate restriction, similarly (L-N) Otx2+/Chx10 cells, early markers of bipolar fated cells. (O-R) An early population of developing photoreceptors (Recoverin+/Crx+) is also present. (S-V) Tuj1+/Brn3+ retinal ganglion cells are present on the inner aspect of the organoids.

Retinal ganglion cells (RGCs, Brn3+ and Tuj1+, figure 3 S-V) were abundant lining primarily the internal perimeter of the organoids. The nuclei of retinal ganglions cells were primarily found in the inner core of the organoid, while their processes could span to the outer aspect of the organoids.

The relative localization of photoreceptors and RGCs corresponds to the lamination seen in the retina *in vivo*. While there isn’t a clear separation between neural retinal progenitors and retinal ganglion cells, the distinct localization of RGC nuclei suggests the possible formation of inner retinal lamination.

Stage 2 retinal organoids are composed of high populations of photoreceptor precursors. During stage 2, the retinal organoids had an increased Crx- and Recoverin positive cells (Figure 4A-D) as well as Otx2 (Figure 4E-G), indicating increase differentiation of photoreceptors. For all these markers, photoreceptors tended to occupy the outer aspect of the organoid, but some were at times observed on the inner portion of the organoid.

**Figure 4.**
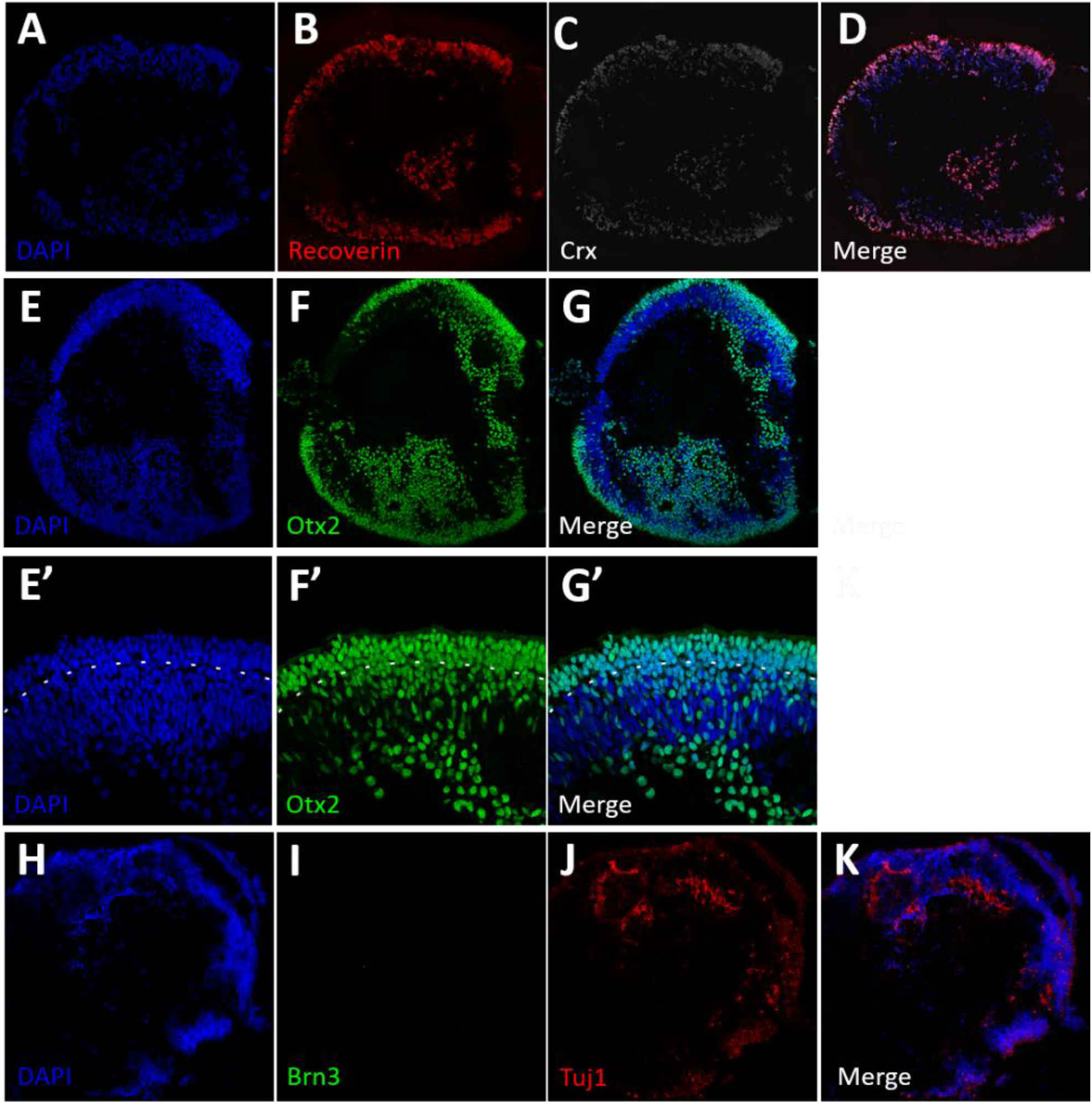
ICC of Stage 2 (80 days): Stage 2 retinal organoids are composed of differentiating photoreceptors and have lost retinal ganglion cells. A-D) Stage 2 retinal organoids have high numbers of developing photoreceptors (Crx+/Recoverin+) and photoreceptors progenitors (Otx2+) (E-G). While a high density of photoreceptors are localized to the outer perimeter, there are also a substantial number of unorganized photoreceptors that are localized in the inner portion of the retinal organoids. E’-G’) Higher magnification images of figure E-G, with a dash line indicating a developing IPL. H-K) Stage 2 retinal organoids are devoid of Brn3+ cells and have sparce Tuj1 staining.

The development of an outer nuclear layer (ONL) can be seen advancing at this stage with a light nuclear stratification (Figure 4E, dashed line). The ONL is 3-5 nuclei in thickness at this stage. Furthermore, outer segments are not present, but the outer perimeter of the organoid is clearly defined. The internal composition of the stage 2 organoids, while not hollow like in stage 1, is disorganized.

By stage 2 RGCs are no longer present (Figure 4H-K). We did not observe BRN3-positive cells (Figure 4I). However, there is some TUJ1-positive signal (Figure 4J,K) that we presume to be neurites of degenerating RGCs that no longer label with Brn3 as the nuclei are probably apoptotic.

Stage 3 retinal organoids have developed outer segment-like structures, which appear as translucent hair-like protrusions covering the majority of the retinal organoids. Similar to what has been observed in human retinal organoids, there is a presence of darkly pigmented cells consistent with RPE (Figure 1C, Stage3 of rhiPSC2431, red dot). Staining for ARR3 (cone arrestin) in Stage 3 demonstrates a high number of photoreceptors in the outer perimeter of these organoids. The development of these Stage 3 characteristics was observed regularly at 105 days, while they were not consistently observed in human retinal organoids regularly until about 200 days. During stage 3, we observed S-Opsins, M/L Opsins, and Rhodopsin-positive cells (Figure 5C, D and H, respectively) localized to the outer segment rather than the cell body, indicating appropriate trafficking of phototransduction machinery in differentiating photoreceptors. While earlier stages had faint separation in the ONL and INL, by this stage we saw that there was a clearer delineation of retinal layers (Figure 5K). The outer layer of stage 3 organoids was composed of an outermost layer of 1-2 cone photoreceptor nuclei (Figure 5G, CRX-positive, M/L Opsin-positive, NRL-negative cells) stacked on a 1-3 rod photoreceptor nuclei layer (Figure 5I, NRL-positive CRX-positive). Cone photoreceptors displayed outer segments that are ARL13B positive (Figure 5R) and have pedicles which are localized with CTBP2 (Figure 5S). The inner layer is composed of both Chx10-positive bipolar cells (Figure 5N) and SOX9-positive Muller Glia (Figure 5X). The inner layer is somewhat less stratified and organized than the outer layer of the organoid at this stage. As we observed in stage 2, there are no longer any retinal ganglion cells (BRN3/TUJ11 double positive cells), but some TUJ1-positive neurites persist (Figure 5U-Y). Similarly, photoreceptors previously observed in the inner aspect of the organoid are for the most part absent; however, occasionally they arrange themselves into rosettes with outer segments pointing toward the center (Figure 5P’-T’). During normal development, newly born photoreceptors also undergo radial somal migration.

### Single cell RNA-seq showed stage-appropriate retinal cell types in rhesus retinal organoids

In order to determine the cell types being produced in rhesus retinal organoids, we used single-cell RNA-seq (scRNA-seq). We chose retinal organoids from early stage 2 and stage 3 for these experiments in order to capture the peaks of differentiation of early and late retinal cell types. The studies of retinal organoids at these stages demonstrated heterogeneity in retinal cell types in the monkey retinal organoids at both of the time points (67 days, and 173 days). A combined total of 23,218 cells were captured and sequenced using 10x Genomics’ single-cell scRNA-seq technology. We aligned the raw sequence data, performed quality control, and normalized the transcriptional profiles using the manufacturer’s Cell Ranger software. The profile data of 19,894 single cells (D67; 11,103 cells and D173; 8,791 cells) were obtained and visualized. A uniform manifold approximation and projection (UMAP) plot was used for the unbiased clustering of single cells from retinal organoids from both time points based on retinal cell-type specific markers (Figure 6A). Combined data after merging and processing two raw count matrices using Seurat are shown (Figure 6B). The proportion of cell populations in clusters (Figure 6C) of the combined data were tabulated, as were dot plots, showing the expression level of known markers for specific retinal cell types (Figure 6D). These experiments confirmed the presence of early and late retinal progenitor populations, RGCs, and amacrine cells in early stage 2 organoids. These data also demonstrate the presence of cone and rod photoreceptors (ratio of 0.49 cone to rod), Muller glia, ON-and OFF-bipolar cells, amacrine cells (AC), and horizonal cells in stage 3 retinal organoids, with an absence of retinal ganglion cells. Cell cycle scoring of retinal organoids (Supplemental Figure 1) as well as violin and feature plots of known retinal cell-type specific markers (Supplemental Figure 2) are also assessed.

**Figure 6.**
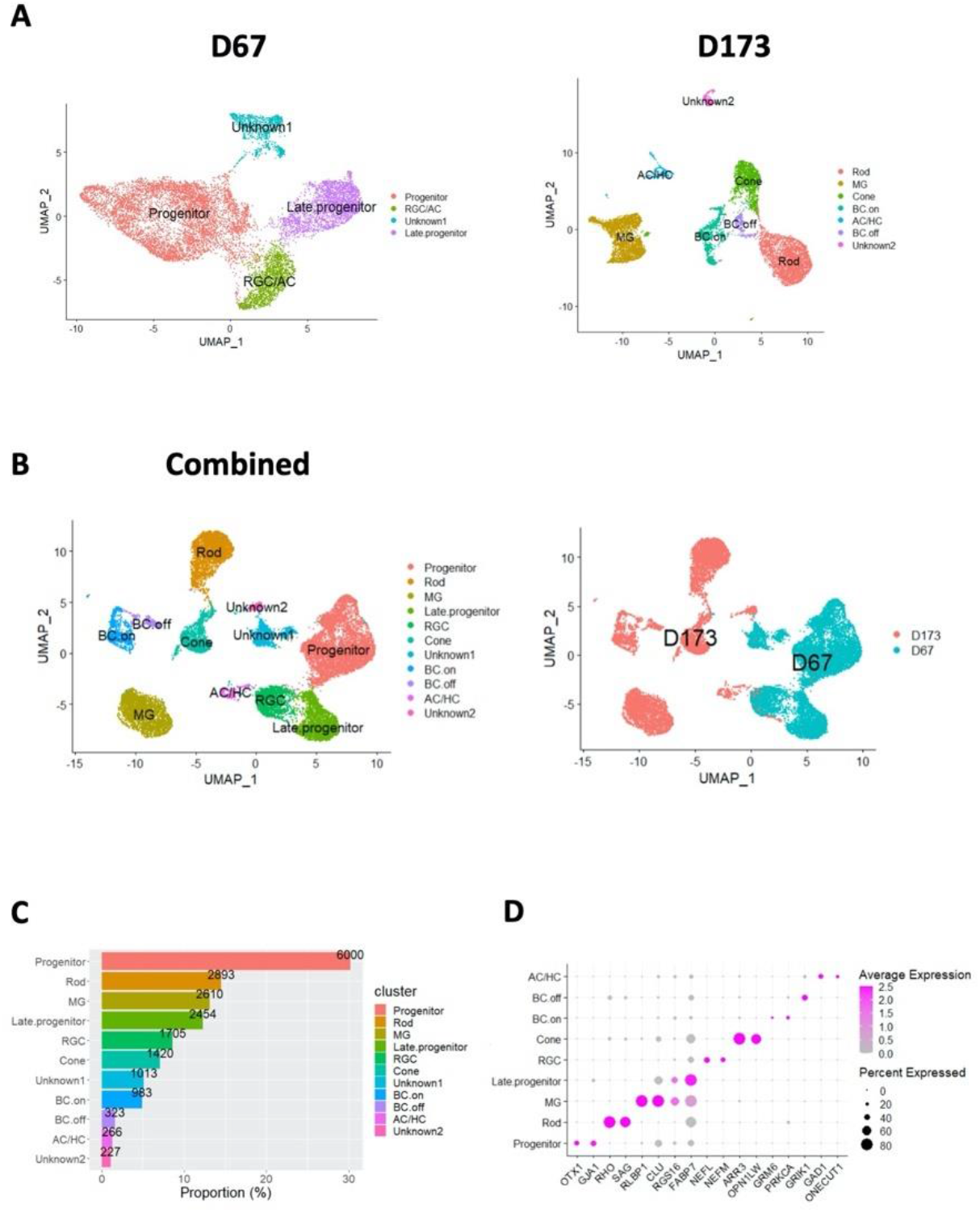
scRNA-seq revealed the heterogeneous cell types in NHP retinal organoids at two different time points (67 days, and 173 days). Total 23,218 cells were captured and sequenced using 10x Genomics’ single-cell RNA-seq (scRNA-seq) technology. After aligning raw sequence data and Quality control and normalization of transcriptional profiles using Cell Ranger software v3.1 (https://www.10xgenomics.com) with default settings and Seurat package (V4.0.1), the profile data of 19,894 single cells were obtained and visualized. (A) Uniform manifold approximation and projection (UMAP) plot was used for the unbiased clustering of single cells from retinal organoids with two different time points (D67; 11,103 cells and D173; 8,791 cells) based on retinal cell specific markers. The standard quality check and workflow for single cell RNA-sequencing analysis were performed using Seurat. Left panel represents output from D67, and right ones for D173. (B) UMAP of the combined data after merging and processing two raw count matrices using Seurat. The UMAPs are visualized by cell type in the left and by each data set in the right. (C) Proportion of cell populations in clusters of the combined data. Numbers on bar indicate the number of cells. (D) Dot plots show the expression level of known cell-type markers for specific cell types: *OTX1* and *GJA1* for progenitor, *RHO* and *SAG* for Rod, *RLBP1* and *CLU* for Muller glia (MG), *RGS16*, and *FABP7* for Late progenitor, *NEFM* and *NEFL* for RGC (Retinal ganglion cell), *ARR3* and *OPN1LW* for Cone, *GRM6* and *PRKCA* for ON Bipolar (BC.on), *GRIK1* for OFF Bipolar (BC.off), *GAD1* for Amacrine cell (AC), *ONECUT1* for Horizonal cell (HC) on the right panel. Cell-types were not assigned for Unknown1 and 2 clusters.

### Rhesus stem cells differentiate precociously and less efficiently compared to human stem cells

We used light microscopy to systematically assign morphological staging levels to retinal organoids based on criteria defined by Capowski et al. 2019. In general, stage 1 retinal organoids developed into stage 2 after a few weeks and stage 3 developed as early as 105 days. The three Rhesus iPSC lines used in this study generally showed similar characteristics in both morphology and timing of differentiation despite somewhat different culturing systems required to maintain them prior to entering the differentiation process. When compared to the H9 human stem cells used for retinal differentiation, we observed rhesus cells passed through the three stages of organoid maturation more quickly than H9 cells (Figure 7A). At day 45, rhesus iPSCs and H9 cells could be seen in stage 1 or 2, and by day 80 most rhesus and human organoids had achieved stage 2 morphology. However, by day 120 virtually all rhesus organoids had developed the outer segment-like projections characteristic of stage 3, while this was never observed in human organoids at this time point. Several rhesus organoids reached stage 3 morphology by day 105. Even at day 150, only a small fraction (∼10%) of human organoids had reached stage 3, which was more commonly attained by day 200. These findings are consistent with the 40% shorter gestational age of the rhesus macaque compared to human. Despite the precocious passage through the stages of retinal organoid differentiation relative to human, Rhesus stem cells underwent self-organization into retinal morphology much less frequently in our protocol. Ninety percent of our differentiation experiments using H9 cells resulted in at least one retinal organoid and were considered successful experiments. However, in rhesus iPSCs, the vast majority of differentiation experiments yielded no retinal organoids, with only 1.9-4.7% of experiments successfully yielding at least one retinal organoid (Figure 7B). Similarly, the proportion of embryoid bodies which adopted retinal morphology was 26.1% in human stem cells, but only 1.4-4.7% in rhesus iPSCs (Figure 7C).

**Figure 7.**
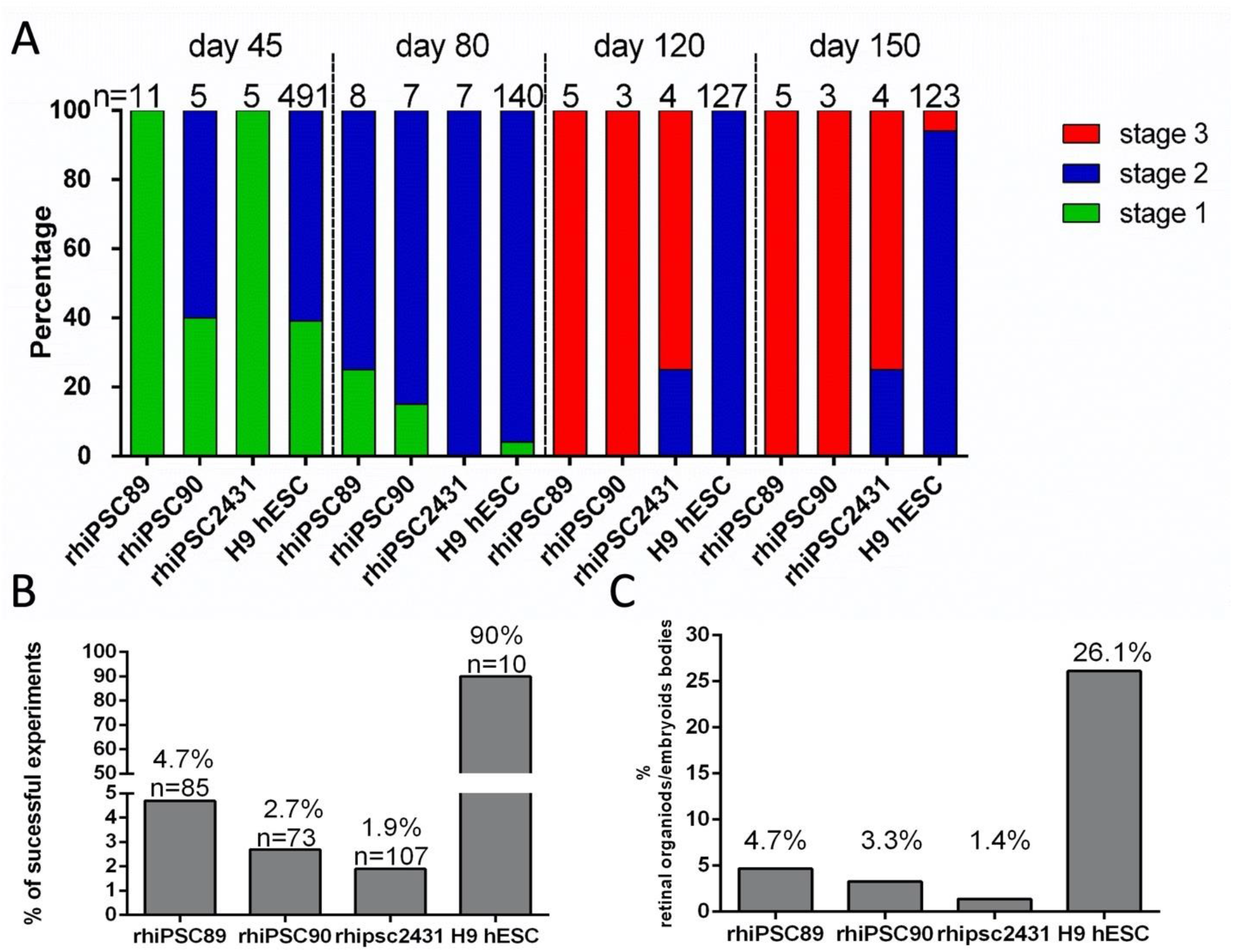
Rhesus iPSCs differentiate precociously but less efficiently than human stem cell derived retinal organoids. (A) Both rhesus and human stem cells generate a significant proportion of stage 1 and 2 retinal organoids at early time points. However, stage 3 morphology marked by outer segment-like projections are seen in the majority of rhesus organoids at day 120, but were never observed in human organoids at this time point. By day 150 approximately 10% of human organoids had achieved stage 3 morphology. (B) The vast majority of differentiation experiments yielded retinal morphology when H9 cells were used, while under 5% of experiments with rhesus cells ever yielded retinal organoids. (C) The proportion of embryoid bodies which adopted retinal morphology was also less than 5% in rhesus cells, while 26.1% of H9 embryoid bodies formed retinal organoids.

## DISCUSSION

Retinal research in NHPs is of particular importance due to the presence of macular and foveal structure, which is required for high acuity vision. The relevance of macular studies to human disease and translation to clinical trials can hardly be overstated. Studies of cell-based therapies ultimately need to be evaluated in the context of allograft transplantation, since human trials will almost certainly involve the use of human cells. Therefore, optimizing the process of cell-based treatments in preclinical models will benefit from using cells of the same species. The value of studying rhesus retinal organoids is that the cells introduced to the NHP host are from the same species. The learnings from this line of experimentation can later be evaluated in the context of human studies once optimized in NHPs. In addition to translational studies, rhesus retinal organoids are useful for studying macular developmental biology and patterning, including cell fate specification of early retinal cell types which make up foveal architecture. The more rapid differentiation quantified in this study emphasizes the potential efficiency of using rhesus iPSCs compared to human stem cells for developmental studies.

The gestational time in humans (280 days, ∼40 weeks) is 40% longer than in Rhesus macaques (166.5 days, ∼23 weeks).^28^ Our hypothesis, based on the difference in gestational age, was that development of retinal organoids and their acquisition of outer segments may occur in an accelerated fashion resulting in relatively precocious differentiation when compared to human derived retinal organoids. Our earliest observation of outer segments was on day 105 in rhesus and day 150 in human organoids. This represents attainment of stage 3 characteristics 30% sooner in Rhesus retinal organoids, which roughly corresponds to the difference in gestational age between these species, especially if taking into account that humans are generally considered premature only prior to 37 weeks.^29^ We have also empirically observed that when embryoid bodies are plated on Matrigel for the self-organization into OVs, rhesus OVs organize slightly sooner, albeit less frequently, than human counterparts. At day 20, clear OVs were visible, while human OVs are just starting to form by this time. As demonstrated in our ICC of each stage, in rhesus macaque derived retinal organoids, there is greater overlap of photoreceptor markers which could not be segregated purely by light microscopy-based binning. This is especially true for stage 1 retinal organoids that demonstrate early photoreceptor markers typically seen in stage 2 human retinal organoids. The formation of an outer nuclear layer which coincides with the loss of retinal ganglion cells are events that occur in stage 2. Studies that seek to enrich for retinal ganglion cells might focus on stage 1 as a termination point to quantify RGC enrichment or to test the efficacy of neuroprotective intervention studies.

Rhesus retinal organoids have the advantage of using a primate cell culture model with much closer genetic and macular structural similarity to humans when compared to mouse retinal organoids. In addition, the development of rhesus retinal organoids is faster in cell culture than humans, which may be advantageous to study mechanistic developmental processes specific to primates. By reducing the time to obtain highly differentiated stage 3 retinal characteristics by 30-40% over human retinal organoids, there may be an advantage of both time to experimental results and reagent expenses. However, given the low efficiency of rhesus organoid production, the differentiation protocol must be optimized for species differences in order to fully implement these advantages.

Various studies have used NHPs as a clinical model for retinal cell replacement, with limited success. A contributing factor to this might be interspecies differences that could be circumvented by using NHP-derived retinal cells for allogeneic transplants, or even autologous. This may be particularly important for pre-clinal trials where both safety and efficacy of delivery of cellular products to the macula would be helpful to demonstrate in NHPs in advance of human clinical trials.

Other uses for rhesus retinal organoids include studies aimed at understanding the developmental processes of the primate retina. There are many questions regarding the genetic patterning of the macula which could be addressed using rhiPSCs as they proceed through the differentiation process faster than human counterparts. Similar questions about cell fate determination of cone photoreceptor subtypes could be addressed. Translational applications of rhesus retinal organoids may help determine if allogenic transplantation of cells to the retina improves integration of donor cells into the host, since human cells may have limited potential to functionally integrate.

Here, we have demonstrated that the derivation of retinal organoids from rhesus macaques is possible and that they closely resemble native retinal tissue. Like what has been observed in human derived retinal organoids, there is high variability between differentiation experiments within the same stem cell lines and across different lines (regardless of iPSCs or hESCs). In this study, we compared three different lines, from which we could obtain retinal organoids, though rhiPSC89 and rhiPSC90 lines produced more retinal organoids that reached stage 3 compared to rhiPSC2431. Rhesus iPSCs are more difficult to culture and may require feeder layers of MEFs since they have a tendency to differentiate prematurely in their absence. The difficulty in maintaining them may overweigh the potential savings in the shorter retinal differentiation time. Optimization of a feeder-free system that is similar to that of human PSCs in an mTSER1 system would be advantageous. Another consideration to take into account is that the differentiation protocol will have to be modified to increase the efficiency of retinal organoid production from rhesus iPSCs. Optimization of the retinal differentiation protocol for rhesus iPSCs is the subject of continued research.

## MATERIALS AND METHODS

### rhiPSC2431 Derivation

Using the Neon transfection system pre-set program #16 (1400V, 20ms, 3 pulses) plasmids used for reprogramming (available from Addgene) were EN2L, ET2K, EM2K, and MIR302, along with EBNA RNA. We transfected 0.5ug of each plasmid into 4 × 10^6 cells per 10 cm Vitronectin-coated dish in DMEM/20%FBS/1X NEAA. After 4-5 days, cells were maintained on vitronectin (5ug/cm^2) in rhesus iPSC media: Essential 8 Flex Medium (Thermofisher) supplemented with 100ug/L rhNodal (R&D #3218-ND), 1.94ug/ml glutathione, 1% Chemically Defined Lipid Concentrate (ThermoFisher), 1% GlutaMAX (ThermoFisher) and 1x Pen/Strep Amphotericin B (Lonza, 17-745E). This line was provided by the Thomson lab and assayed for standard pluripotency markers and had a normal karyotype (supplemental figure 1). rhIPSC89 and rhiPSC90 were maintained on MEFs (50K/cm^2) (Thermofisher, A34180) in hESC media: 80% DMEM/F12, 20% KOSR, 10ng/ml bFGF, 1mM L-glutamine, 0.1mM B-Me, 1% NEAA and 1x Pen/Strep Amphotericin B (Lonza, 17-745E). For feeder-free culture, healthy iPSC colonies were manually picked and plated on reduced growth factor Matrigel plates (50ug/cm^2) in MEF conditioned hESC media supplemented with bFGF prior to use. Human Embryonic stem cell line H9 (WiCell) and CRX +/tdtomato were maintained in reduced growth factor Matrigel (50ug/cm^2) in mTeSR1 plus with 1x Pen/Strep Amphotericin B (Lonza, 17-745E).

### Retinal Differentiation

Spontaneously differentiated cells or colonies with unhealthy morphology were scraped off before beginning of differentiation. Colonies were briefly treated with dispase (1U/ml DMEM/F12) until the edges were slightly curled, at which point the dispase was removed and the colonies were washed with HBSS. Neural induction media (DMEM:F12 1:1, 1% N2 supplement, 1× MEM nonessential amino acids (MEM NEAA), 1× GlutaMAX (Thermo Fisher) and 2 mg/ml heparin (Sigma)) was added and the colonies were scraped off and triturated. The fragmented colonies were then plated in low adhesion plates (coated with poly-HEMA (8mg/cm^2)) and corresponding iPSC media was added to achieve a 3:1 PSC media/neural induction media (NIM). The embryoid bodies were sequentially weaned into neural induction media with 50% media changes replaced by fresh NIM. On day 5, a 50% NIM change was done and on day 6, the EB were treated with 1.5nM recombinant human BMP4 (PeproTech, 10-05ET) in fresh NIM. On day 7, the EBs were plated on reduced growth factor Matrigel (50EBs/well of a 6-well plate; 30EBs/cm^2). On day 9 and 12, 50% media was replaced with fresh NIM. On day 15, an additional 50% NIM was added and the following day, the media was replaced entirely with fresh retinal differentiation media (RDM) (DMEM:F12 3:1, 2% B27 supplement, MEM NEAA, 1× antibiotic, antimycotic (Thermo Fisher) and 1× GlutaMAX)). Retinal Differentiation media was changed every other day until day 30. The 3 dimensional structures with lamination like and optic vesicle like appearance—retinal organoids--were picked off the plate and resuspended on polyHEMA coated plates in 3D-RDM (DMEM:F12 3:1, 2% B27 supplement, 1× MEM NEAA, 1× antibiotic, anti-mycotic, and 1× GlutaMAX with 5% FBS, 100 µM taurine, 1:1000 chemically defined lipid supplement (11905031, Thermo Fisher)) supplemented with 1uM of all-trans-retinoic acid until day 100. 3D-RDM either with or without RA supplementation was changed twice per week.

### Immunocytochemistry

Retinal organoids were fixed with 4% PFA on ice for 20min and then washed three times with DPBS. The retinal organoids were then equilibrated in 15% sucrose and 30% sucrose until they sunk. The retinal organoids were then flash frozen in dry ice-ethanol bath in Tissue-Plus OCT compound. Subsequently, the blocks were sectioned at 10um and blocked in blocking solution (4% BSA and 0.5% Triton X-100 in PBS) for an hour at room temperature. Primary antibodies were then incubated at 4C overnight. Excess primary antibody was washed off with three washes of PBS. Alexa Fluor conjugated secondary antibodies matching the primary antibody host were incubated for an hour followed by 5min incubation in DAPI, similarly, the slides were washed three times, one minute each. Finally, the slides were cover slipped with FluorSave Reagent (Millipore, 345789). Description of primary antibodies and secondary antibodies are included in supplementary table 1. The samples were then imaged using an Olympus FluoView FV1000 spectral confocal microscope.

**Table 1:** Primary antibodies.

| Antibody | Manufacturer | catalog number | dilution |
| --- | --- | --- | --- |
| ARL13b | Protein Tech | 17711-1-AP | 1:500 |
| Bassoon | Enzo Life Sciences | ADI-VAM-PS003-F | 1:200 |
| Blimp-1/PRDI-BF1 | Cell Signaling Technology | C14A4 | 1:100 |
| Blue (S) Opsin | EMD Millipore | AB5407 | 1:1000 |
| Brn-3 | Santa Cruz Biotechnology | sc-6026 | 1:300 |
| Calretinin (CaR) | Swant | CG1 | 1:400 |
| Chx10 | Santa Cruz Biotechnology | sc-365519 | 1:100 |
| Cone arrestin (ARR3) | Novus Biologicals | NBP1-37003 | 1:500 |
| Cone arrestin (ARR3) | LSBio | LS-C368677-200 | 1:500 |
| CRX | Abnova | H0001406-M02 | 1:1000 |
| C-Terminal Binding Protein-2 | BD Biosciences | 612044 | 1:300 |
| Ki67 | ThermoFisher Scientific | RM-91060-S | 1:200 |
| LHX2 | Santa Cruz Biotechnology | sc-517243 | 1:100 |
| NR2E3/RNR | Abcam | ab172542 | 1:300 |
| NRL | R&D Systems | AF2945 | 1:300 |
| OTX2 | R&D Systems | AF1979 | 1:500 |
| Pax-6 | BioLegend | 901302 | 1:100 |
| Pericentrin (PCN) | Abcam | ab28144 | 1:500 |
| PKCα | Sigma | P4334 | 1:50 |
| Recoverin (RCVN) | EMD Millipore | AB5585 | 1:1000 |
| Red/Green (M/L) Opsin | EMD Millipore | AB5405 | 1:1000 |
| Rhodopsin (RHO) | EMD Millipore | MABN15 | 1:500 |
| Rx | Takara Bio | M229 | 1:400 |
| Sox9 | EMD Millipore | AB5535 | 1:1000 |
| Synuclein $\gamma$ (SNCG) | Abnova | H00006623-M01A | 1:500 |
| Tubulin β3 (TUBB3) (Tuj1) | BioLegend | 801202 | 1:500 |
| VGLUT1 | NeuroMab | 75-066 | 1:200 |

### TEM

Organoids were fixed in 3% glutaraldehyde and 1% paraformaldehyde in 0.08 M sodium acodylate buffer (all from Electron Microscopy Sciences) overnight with gentle rocking at 4°C, washed with 0.1 M cacodylate buffer, and post-fixed in 1% osmium tetroxide for 2 h at RT. The organoids were then dehydrated in a graded ethanol series, further dehydrated in propylene oxide and embedded in Epon epoxy resin. Semi-thin (1 μm) and ultra-thin sections were cut with a Leica EM UC6 ultramicrotome and the latter were collected on pioloform-coated (Ted Pella, 19244) one-hole slot grids. Sections were contrasted with Reynolds lead citrate and 8% uranyl acetate in 50% ethanol and imaged on a Philips CM120 electron microscope equipped with an AMT BioSprint side-mounted digital camera and AMT Capture Engine software.

### Single cell sequencing and data analysis

Single cell cDNA library preparation and sequencing were performed following manufacturer’s protocols (https://www.10xgenomics.com). Single cell suspension at 1000 cells/µl in PBS will be loaded on a Chromium controller to generate single cell GEMS (Gel Beads-In-EMulsions). The scRNA-Seq library is prepared with chromium single cell 3’ reagent kit v3 (10x Genomics). Cell Ranger software v3.1 (https://www.10xgenomics.com) with default settings was used for alignment, barcode assignment and UMI counting of the raw sequencing data with genome reference hg19. After generating UMI count profile, we applied Seurat 4.0 (https://satijalab.org/seurat) for quality control and downstream analysis.

For Quanlity control we excluded genes detected in less than 3 cells, and cells were filtered out if UMI counts are less than bottom 3 % and greater than top 1% of total quantile. We removed cell cycle effects by regressing out cell cycle scores during data scaling using of all signals associated with cell cycle using “CellCycleScoring” function in Seurat. Next, a global-scaling normalization method ‘LogNormalize’ in Seurat is employed to normalize the gene expression measurements for each cell by the total expression, then the result is multiplied by a scale factor (10,000 by default), and log-transformed. We selected variable genes and computed principle components for dimensional reduction of UMAP with default parameters of Seurat. Next, we performed the clustering using ‘FindClusters’ in Seurat to identify sub-cell type clusters. Top 20 principle components were used with 0.1 resolution and the sub-populations of corneal cells are visualized using UMAP. To identify differentially expressed marker genes in each cluster, we used the ‘FindAllMarkers’ function based on the Wilcoxon rank-sum test in Seurat with default parameters, and then cell types were assigned using known cell-type markers in Fig2a. Top 10 DEGs were visualized with a heatmap using Seurat. We also performed cell cycle analysis to estimate cell cycle status of each cell using cell cycle markers using Seurat.

For trajectory analysis we used Monocle 3 to infer the cell differentiation trajectories with default parameters (https://cole-trapnell-lab.github.io/monocle3/). This method places the cells along a trajectory corresponding to a biological process (in our case, cell differentiation) by taking advantage of an individual cell’s asynchronous progression under an unsupervised framework.

## FIGURES AND LEGENDS

**Sfig1.**
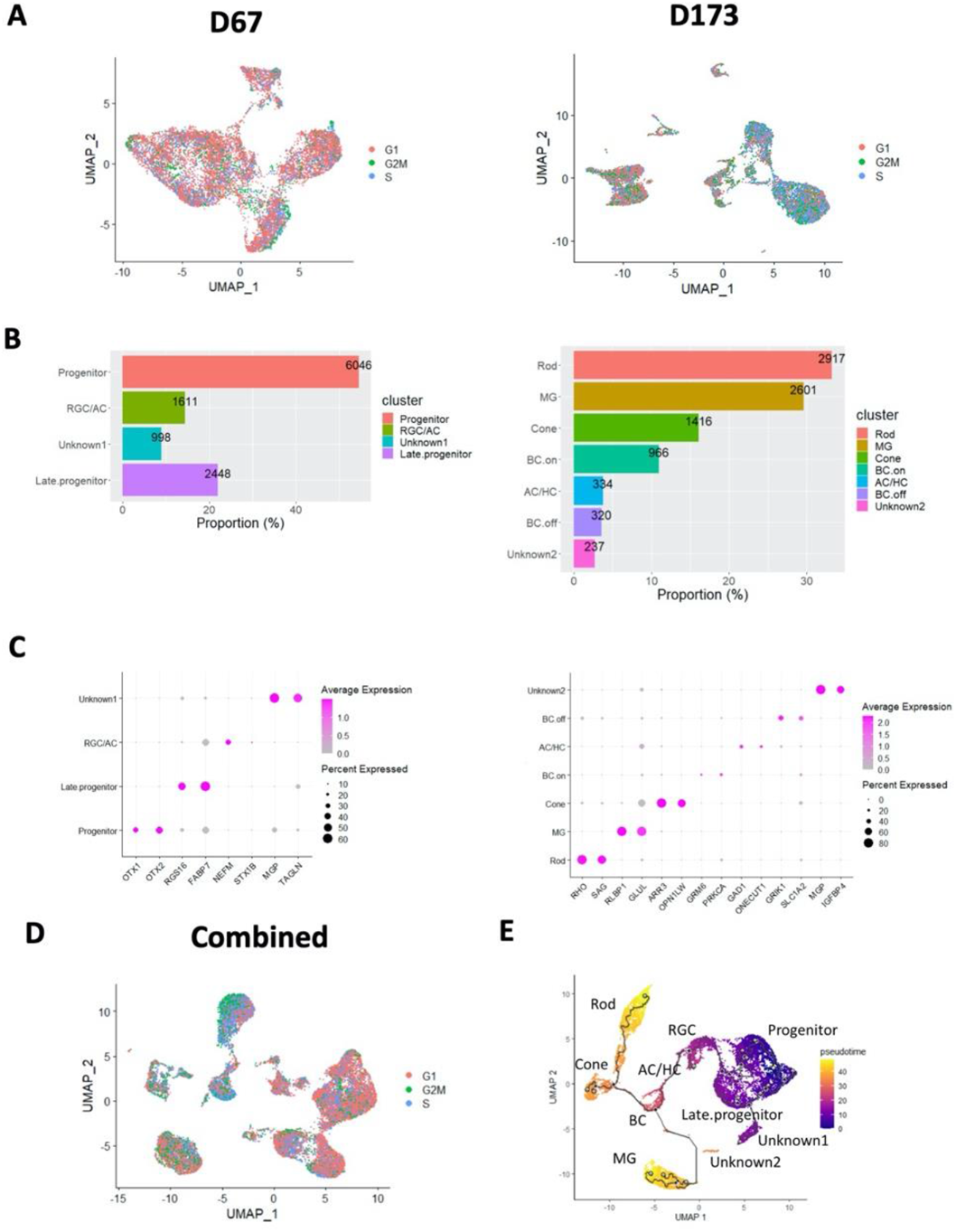
UMAP of organoid data sets with cell cycle scoring. (A) The cell cycle stages for single cells per cluster in D67 (left) and D173(right) data, were predicted by subpopulation using the Cell Cycle Scoring function of the Seurat package and regressed out all cell cycle effects. Each stage represents in color: Red for S1, green for G2/M, and blue for S. (B) Cell populations in D67 (left) and D173(right) data (C) Dot plots show the expression level of known cell-type markers for specific cell types in D67 (left) and D173(right) data : *OTX1* and *OTX2* for progenitor, *SPP1* for aRGC, *NEFM* and *NEFL* for RGC (Retinal ganglion cell), *STX1B* for Amacrine cell (AC) on the left panel; *RHO* and *SAG* for Rod, *RLBP1* and *CLU* for Muller glia (MG), *ARR3* and *OPN1LW* for Cone, *GRM6* and *PRKCA* for ON Bipolar (BC.on), *GAD1* for Amacrine cell (AC), *ONECUT1* for Horizonal cell (HC), and *GRIK1* and *SLC1A2* for OFF Bipolar (BC.off) on the right panel. Expression of known cell-type markers in D67 (left) and D173(right) data. Cell-types were not assigned for Unknown1 and 2 clusters. (D) The cell cycle stages in the combined data. (E) Pseudotemporal trajectory plot of the combined data with cell types. The single-cell trajectory was predicted by Monocle 3 and visualized by UMAP. Cells are ordered in pseudotime colored in a gradient from purple to yellow.

**Sfig2.**
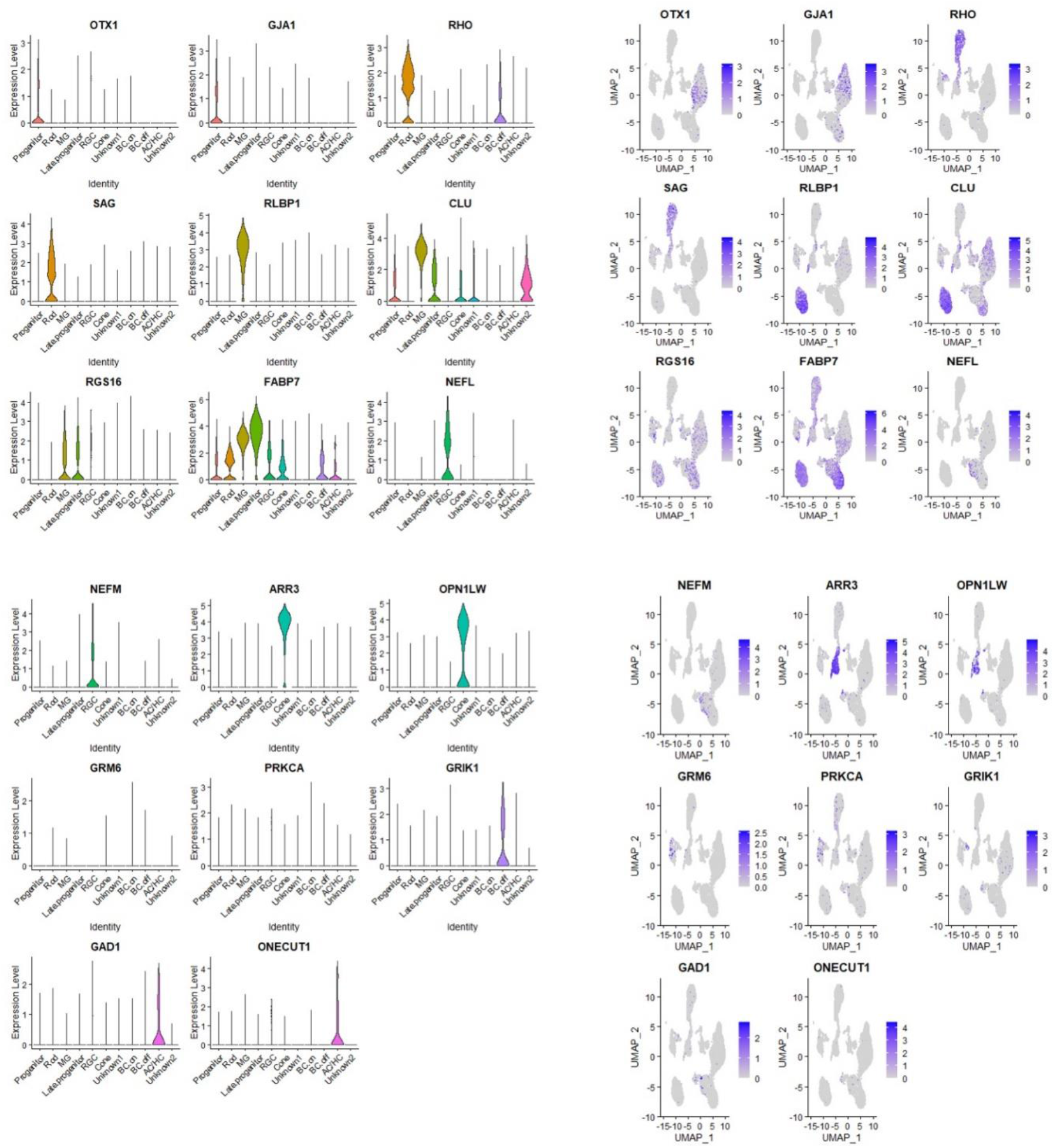
Violin and feature plots of known cell-type markers in the combined data. (A) Violin and (B) feature plots for known cell-type markers were used for classifying individual cell type in the combined data such as OTX1 and GJA1 for progenitor, *RHO* and *SAG* for Rod, *RLBP1* and *CLU* for Muller glia (MG), *RGS16* and *FABP7* for Late progenitor, *NEFL* and *NEFL* for RGC, ARR3 and OPN1LW for Cone, GRM6 and PRKCA for ON Bipolar (BC.on), *GRIK1* and *SLC1A2* for OFF Bipolar (BC.off), *GAD1* for Amacrine cell (AC), and *ONECUT1* for Horizonal cell (HC). Cell-types for two clusters were not assigned as Unknown1 and Unknown2.

